# ZBP1 Senses Splicing Aberration through Z-RNA to promote Cell Death

**DOI:** 10.1101/2025.03.24.645142

**Authors:** Zhang-Hua Yang, Puqi Wu, Bo-Xin Zhang, Cong-Rong Yang, Jia Huang, Lei Wu, Shuang-Hui Guo, Yuenan Zhou, Yuanhui Mao, Yafei Yin, Rongbin Zhou, Hanming Shen, Zhi-Yu Cai, Wei Mo

**Author notes:** These authors contributed equally to this work (Zhang-Hua Yang, Puqi Wu). Corresponding author. (W.M.); (Z.-Y.C.); (Z.-H.Y.).

## Abstract

RNA splicing, a highly regulated process performed by the spliceosome, is crucial for eukaryotic gene expression and cellular function. Numerous cellular stresses including oncogenic insults dysregulate RNA splicing, often provoking inflammatory responses and cell death. However, the molecular signal produced by spliceosome aberration and how cell sensed and respond to it remain elusive. Here we show that spliceosome inhibition induces the widespread formation of unique, left-handed nucleic acids, Z-form nucleic acids (Z-NAs) from the nucleus. These Z-NAs were double-stranded RNA (dsRNA), rather than DNA-RNA hybrids, that were predominantly derived from transcripts of mis-spliced intronic RNA. Spliceosome inhibition induced the egress of these Z-RNA from the nucleus to the cytoplasm in an active manner. Sensing of the accumulation of Z-RNA in the cytosol by the host sensor ZBP1 triggered cell death, mainly for RIPK3-MLKL dependent necroptosis. Collectively, these findings delineate a previously uncharacterized mechanism in which Z-NA sensing by ZBP1 responds to global aberrations of RNA splicing to trigger inflammatory cell death.

## Introduction

RNA splicing is a highly regulated process that occurs inside the nucleus, carried out by the spliceosome—a large complex made up of both RNA and protein components—along with additional regulatory splicing factor proteins that fine-tune its activity^1-3^. Numerous examples of alternative splicing events are crucial for cell identity, pluripotency, and organismal physiology, and they can contribute to various pathologies^3-7^. Dysregulated alternative splicing has emerged as a significant mechanism in the pathogenesis of diverse diseases, including cancer and neurodegenerative disorders^7-10^.

Mutations or expression changes affecting components of the splicing machinery or splicing factors have been intensely studied for their roles in cancer initiation and progression^11^. Recurrent somatic mutations in key spliceosome components, such as SF3B1, SRSF2, U2AF1, and ZRSR2, frequently occur in hematological malignancies^9, 10^. Cancer cells, particularly those with spliceosome mutations, exhibit greater sensitivity to compounds that inhibit the spliceosome compared to non-cancer cells^12^. This has led to the development of therapeutic agents targeting the spliceosome^13^.

It is hypothesized that spliceosome-targeted therapies (STTs) induce cancer cell death by affecting common splicing events^14^. Indeed, mutations in major splicing factors have been shown to alter numerous splicing events in cellular and animal models. While some mis-spliced candidates have been linked to therapeutic responses to STTs, each splicing factor mutation appears to cause a distinct set of splicing changes in both cellular models and patients. Additionally, different structural variants of spliceosome-targeting compounds lead to alternative splicing changes that only partially overlap^15, 16^. These findings suggest that a generalized mechanism, in addition to or independent of splicing changes, may contribute to cancer cell cytotoxicity following global spliceosome inhibition. Therefore, it is crucial to molecularly define the cell death pathways activated by and coordinating cell-fate decision-making in response to spliceosome inhibition to understand the molecular mechanisms by which STTs inhibit tumorigenesis.

Previous studies have reported that RNA splicing perturbation promotes R-loops accumulation that detected by the S9.6 antibody^17-20^. However, it is well-documented that besides recognized RNA:DNA hybrids, the S9.6 antibody also has significant affinity for double-stranded RNA (dsRNA)^21^. A recent study has showed that deletion of individual of dsRNA sensor as well as their signaling integrator MAVS only partially mitigated the cell death induced by spliceosome inhibition^14^. The bona-fide sensor of these R-loops or dsRNA and their effects on the cell fate under spliceosome inhibition remains unclear.

ZBP1 is a Z-form nucleic acids sensor, which connects nucleic acids sensing to programmed cell death. The recognition of exogenous or endogenous Z-nucleic acids by ZBP1 triggers its activation, leading to the recruitment of RIPK3, which induces parallel pathways of caspase-8-mediated apoptosis and MLKL-mediated necroptosis^22-27^. Increasing examples of abnormal nucleic acids produced in cells under various cellular stresses, including epigenetic dysregulation and replicative crisis, have been shown to be sensed by ZBP1 to determine cell fate, indicating that ZBP1 is a surveillance mechanism for the homeostasis of cellular nucleic acids^22, 28, 29^.

Here we revealed that the spliceosome inhibition caused widespread accumulation of intron-containing transcripts that produced Z-form double stranded RNA (Z-RNA) in cells. This accumulated Z-RNA binds to the pattern recognition receptors ZBP1, activating RIPK3 and inducing necroptosis and apoptosis. These findings establish Z-RNA as an immunogenic species, similar as DAMPs, that aberrantly accumulate in the cell following spliceosome inhibition, linking splicing aberrations to cell death through the innate immune response. Aberrant RNA splicing and subsequent innate immune activation may contribute to various diseases, including neurodegeneration and cancer.

### Spliceosome inhibition triggers ZBP1-mediated cell death

Spliceosome inhibition leads to the activation of host defense^11, 14^. ZBP1 is an interferon-stimulated gene (ISG) responding to host defense. Our previous study revealed that the alternative splicing of *Zbp1* represents autogenic inhibition for regulating ZBP1 signaling^30^. To assess whether regulation of splicing could modulate the strength and kinetics of ZBP1-dependent cell death, we used pladienolide B (PladB), a widely used spliceosome inhibitor, to induce splicing dysfunction in MEF cells. qPCR analysis revealed that PladB treatment induced rapid pre-mRNA retention in MEF cells (Figure S1A). Immunoblotting analysis revealed that the expression of the long and especially the short isoform of mouse ZBP1 induced by IFN was dramatically decreased by PladB treatment (Figure S1B). Despite the reduced ZBP1 expression by PladB treatment, we observed that PladB treatment induced obvious lytic cell death in IFN-priming wildtype (WT) MEF cells (Figure 1A) in a ZBP1-dependent manner. Deletion of *Zbp1* markedly prevented PladB-induced cell death (Figure 1A). Immunoblot analysis showed that PladB treatment induced both necroptosis (phosphorylation of MLKL) and apoptosis (proteolytic activation of caspase-3) in IFN-priming WT MEFs (Figure 1B). And both signals were entirely dependent on ZBP1 as *Zbp1* deficiency (*Zbp1* KO) completely prevented these signals in the cells (Figure 1B). Consistent with the inhibitory role of caspase-8 in ZBP1-mediated cell death, addition of pan-caspase inhibitor zVAD, significantly accelerated PladB-induced ZBP1-dependent cell death (Figure 1A). These results are not unique to MEF cells, as treatment of L929 with PladB resulted in similar ZBP1-dependent cell death (Figure 1C, D). These data demonstrated that spliceosome inhibition by PladB induced ZBP1-mediated necroptosis and apoptosis. Of note, the cell death-induced by PladB is independent of the autocrine TNF signaling, as deletion of *Tnfr1* had no effect on the PladB-induced cell death (Figure S1C).

**Figure 1.**
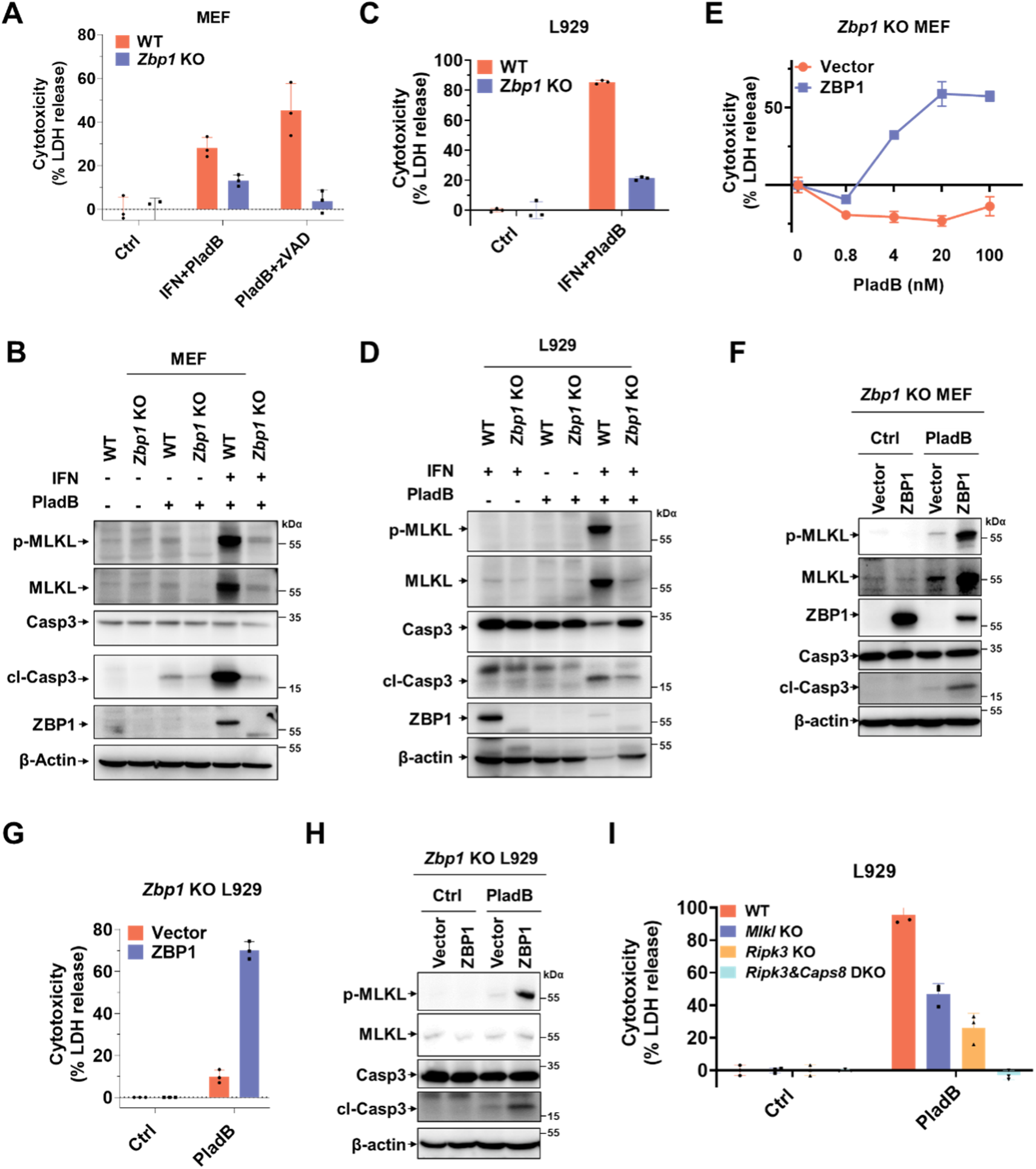
Inhibition of spliceosome initiates ZBP1-dependent cell death. **(A)** Quantitation of cell death by measuring LDH release from MEF cells treated with PladB (100 nM) plus inteferon (IFN) (50 ng/ml, priming for 24 h) or Plad B (100 nM) plus zVAD (20 μM) for 24 h. **(B)** Immunoblot analysis of cell lysates from wildtype (WT) and *Zbp1* knockout (KO) MEFs treated as indicated for 12 h. Phosphorylated MLKL (p-MLKL) was detected in the RIPA insoluble fraction of cell lysates. Identical methods were used in later experiments unless otherwise stated. cl-Casp3: cleaved caspase-3. **(C)** Quantitation of cell death by measuring LDH release from L929 cells treated with PladB (100 nM) plus IFN (50 ng/ml, priming for 24 h) for 24 h. **(D)** Immunoblot analysis of cell lysates from WT and *Zbp1* KO L929 cells treated as indicated for 12 h. p-MLKL was detected in the RIPA insoluble fraction of cell lysates. cl-Casp3: cleaved caspase-3. **(E)** *Zbp1* KO MEFs reconstituted with empty vector or full-length murine ZBP1 were treated as indicated for 12 h and the cell death was measured by LDH release. **(F)** *Zbp1* KO MEFs reconstituted with empty vector or full-length murine ZBP1 were treated as indicated for 12 h and the cell lysates were analyzed by immunoblot as indicated. **(G)** *Zbp1* KO L929 cells reconstituted with empty vector or full-length murine ZBP1 were treated as indicated for 24 h and the cell death was measured by LDH release. **(H)** *Zbp1* KO L929 cells reconstituted with empty vector or full-length murine ZBP1 were treated as indicated for 24 h and the cell lysates were analyzed by immunoblot as indicated. **(I)** IFN-priming WT, *Mlkl* KO, *Ripk3* KO or *Ripk3&Casp8* double knockout (DKO) L929 cells were treated with PladB for 24 h and the cell death was measured by LDH release. Data are representative of three independent experiments. Error bars represent mean ± SD. *p<0.05, **p<0.01, ***p<0.001, ****p<0.0001.

To further confirm the indispensable role of ZBP1 in mediating PladB-induced cell death, we stably reconstituted the expression of ZBP1 in *Zbp1* KO MEF cells, to bypass the extra effects of IFN-priming. As shown in Figure 1E, restoration of ZBP1 expression dramatically sensitized the cells to PladB-induced cell death. Immunoblot analysis confirmed the robust occurrence of both necroptosis and apoptosis (Figure 1F). As expected, this cell death was independent of IFN priming (Figure S1D), confirming that the role of IFN-treatment is to upregulate the expression of ZBP1. Consistent with MEF, similar results have been obtained in *Zbp1* KO L929 reconstituted with ZBP1 (Figure 1G, H). These results indicated that ZBP1 sensitizes cells to PladB-induced cell death.

Next, we sought to understand whether ZBP1 commonly mediated cell death by spliceosome inhibitions. Treatment of ZBP1-reconstituted MEF cells using Madarsin— a structurally distinct SF3B1 inhibitor—also induced ZBP1-dependent cell death (Figure S1E). Immunoblot analysis of the protein extracts from Madarsin-treated MEFs revealed phosphorylation of MLKL, as well as cleavage of caspase-3 in a ZBP1-dependent manner (Figure S1F). Another broad-spectrum spliceosome inhibitor— isoginkgetin, which prevents recruitment of the U4/U5/U6 tri-snRNP and leads to stalling at the pre-spliceosomal A complex, phenocopied the effects of PladB and Madarsin (Figure S1G, H). Together, these results demonstrated that spliceosome inhibition triggers ZBP1-mediated cell death.

We next aimed to understand the pathways downstream of ZBP1 that are activated by spliceosome inhibition. We observed that deletion of *Ripk3* and *Mlkl* substantially, though not entirely, rescued the cell death induced by PladB, while the combined deletion of *Ripk3* and *Casp8* could completely blocked PladB-induced cell death (Figure 1I). In line with this, combined pharmacological inhibition of RIPK3 and caspases strongly reduced the PladB-induced death of ZBP1-reconstitution MEFs (Figure S1I). Together, these results demonstrate that ZBP1 activates both apoptosis and necroptosis following spliceosome inhibition, similar to what have been observed in other scenarios of ZBP1 activation, such as after *Setdb1* loss and during influenza A infection.

### Zα-dependent sensing of Z-NAs induced ZBP1-mediated cell death upon spliceosome inhibition

ZBP1 contains two N-terminal Zα domains, Zα1 and Zα2, which are responsible for ligand sensing, followed by two receptor-interacting protein homotypic interaction motif (RHIM) domains that have the capacity to recruit other RHIM proteins, like RIPK3^31, 32^. To determine whether the cell death induced by spliceosome inhibition requires Zα-dependent sensing of endogenous ligands and RHIM-dependent signaling, we mutated the two Zα domains (Zα1 and Zα2 double mutant [ZαDM]) and the RHIM domain (RHIMmut), and reconstituted *Zbp1* KO MEF cells with these mutants. Immunoblot analysis confirmed that the mutant ZBP1 proteins were expressed similarly to WT ZBP1 (Figure 2A). Indeed, PladB treatment induced cell death in cells that expressed with WT-ZBP1, but not in those expressing ZBP1 with mutations of the two Zα domains or the RHIM domain (Figure 2A). Consistent with the cell death results, the signals of phosphorylation of MLKL and cleavage of caspase-3 induced by PladB were nearly completely blocked in cells expressed with mutant ZBP1 (Figure 2B). Furthermore, we obtained similar results in reconstituted L929 cells (Figure 2C, D). In line with the cell death phenotype, the cell death induced by other spliceosome inhibitors, Madarsin and isoginkgetin, were also dependent the Zα and RHIM domains of ZBP1 (Figure 2E). Collectively, Zα-dependent sensing of endogenous ligands and RHIM-mediated signaling by ZBP1 causes cell death in cells with spliceosome inhibition.

**Figure 2.**
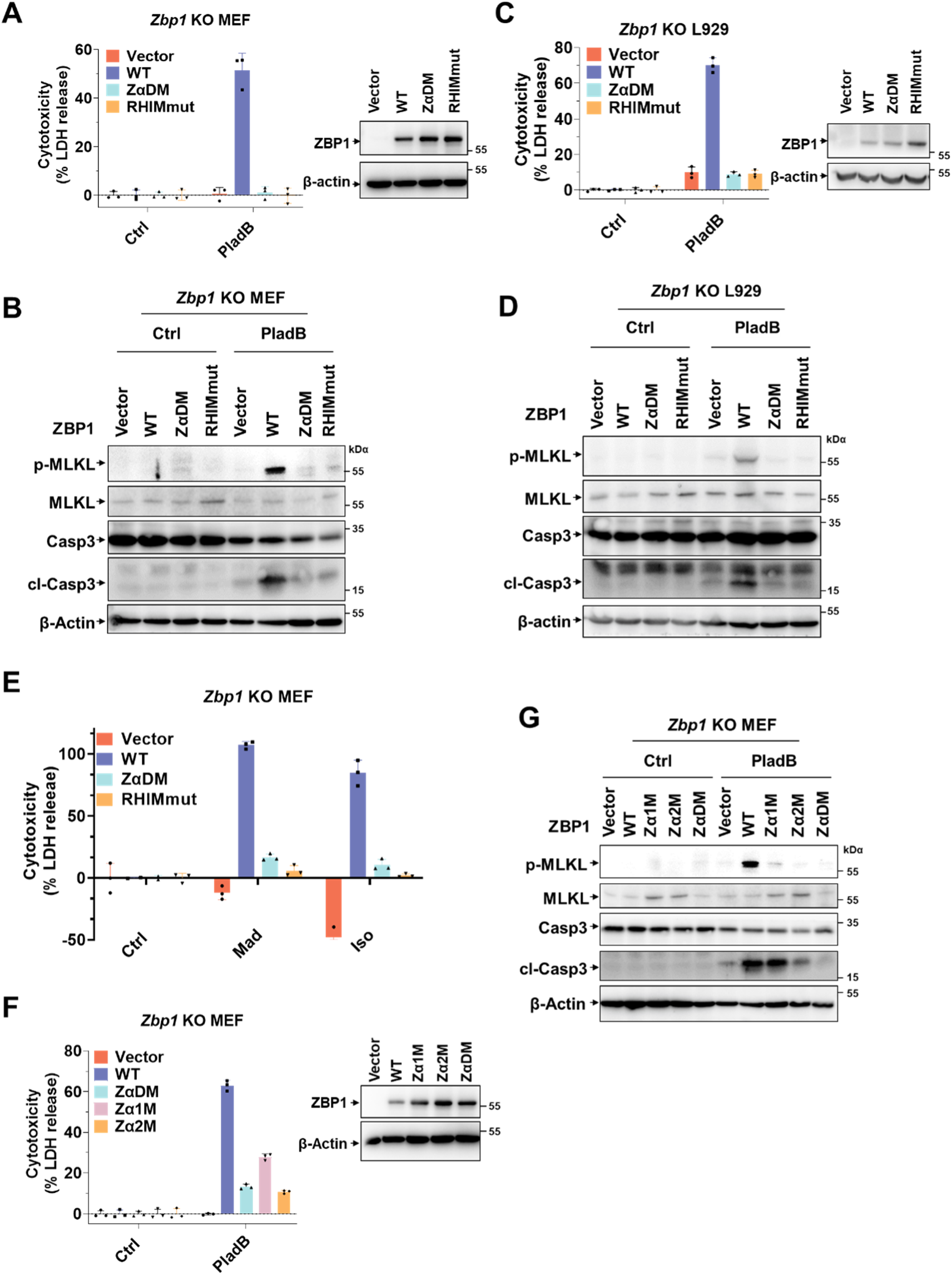
Zα and RHIM domains of ZBP1 are required for cell death induced by spliceosome inhibition. **(A)** *Zbp1* KO MEFs reconstituted with empty vector, WT ZBP1 or the indicated ZBP1 mutants were treated with PladB for 24 h. the cell death was measured by LDH release and the expression levels of ZBP1 were analyzed by immunoblot. **(B)** The same cells as in (**A**) were treated with PladB for 12 h and the cell lysates were analyzed by immunoblot as indicated. **(C)** *Zbp1* KO L929 cells reconstituted with empty vector, WT ZBP1 or the indicated ZBP1 mutants were treated with PladB for 24 h. the cell death was measured by LDH release and the expression levels of ZBP1 were analyzed by immunoblot. **(D)** The same cells as in (**C**) were treated with PladB for 24 h and the cell lysates were analyzed by immunoblot as indicated. **(E)** *Zbp1* KO MEFs reconstituted with empty vector, WT ZBP1 or the indicated ZBP1 mutants were treated as indicated for 12 h. the cell death was measured by LDH release and the expression levels of ZBP1 were analyzed by immunoblot. **(F)** *Zbp1* KO MEFs reconstituted with empty vector, WT ZBP1 or the indicated ZBP1 mutants were treated with PladB for 24 h. the cell death was measured by LDH release and the expression levels of ZBP1 were analyzed by immunoblot. **(G)** The same cells as in (**F**) were treated with PladB for 12 h and the cell lysates were analyzed by immunoblot as indicated. Data are representative of three independent experiments. Error bars represent mean ± SD. *p<0.05, **p<0.01, ***p<0.001, ****p<0.0001.

ZBP1 contains two Zα domains that selectively bind left-handed double-helical “Z-form” nucleic acids. To further figure out which Zα domain is responsible for the sensing of endogenous ligands, we reconstituted the single Zα1 mutant or Zα2 mutant ZBP1 in *Zbp1* KO MEF cells, respectively (Figure 2F). As shown by the fact that mutation of Zα2 abolished the function of ZBP1, similar to ZαDM (Figure 2G). Notably, the Zα1 mutant of ZBP1 also substantially, albeit not completely, rescued the cell death induced by PladB. These data indicated that Z-nucleic-acid binding to ZBP1 is pivotal in activating ZBP1 upon splicing inhibition, with the Zα2 domain being particularly essential.

### Spliceosome inhibition caused the accumulation of Z-RNA from intronic regions

Previous studies have showed that ZBP1 senses viral or endogenous Z-RNA/Z-DNA through its Zα domains, and binding of Z-NA activates ZBP1. We next sought to understand how spliceosome inhibition produced ligand for the activation of ZBP1 through its Zα domain. Cells were examined for the presence of Z-NA using an antibody (clone Z22) that could specifically recognize both DNA and RNA in Z-form. Treating MEF cells with PladB manifested a predominantly nuclear signal when stained with the Z22 antibody, detectable by 4 hours after PladB treatment, gradually increasing in intensity over 12 hours (Figure 3A). Of note, the Z22 signals increased and spread to the cytoplasm from 8 hours post-PladB treatment (Figure 3A). Immunofluorescence staining using a dsRNA-specific antibody (J2 antibody) revealed that most of the Z22 signals were overlapping with J2 signal, with kinetics of induction largely paralleling that of Z22 (Figure 3A). Furthermore, the signal produced by the Z22 antibody in PladB-treated MEFs was sensitive to RNase A, as well as dsRNA-specific RNase III treatment, but not to DNase I (Figure 3B, C), which suggests that it originated from the accumulation of endogenous dsRNAs rather than DNA. Since previous studies have reported that PladB treatment could induce the formation of R-loops, we examined whether the Z-NAs induced by PladB involved RNA:DNA hybrids^33, 34^. RNase H treatment, which specifically digested RNA:DNA hybrids (Figure S2C), had no impact on the Z-NA signals produced by PladB treatment (Figure 3D). Together, these data indicate that the Z-NA produced PladB is dsRNA, rather than RNA:DNA hybrids. Consistent with the cell death data, other spliceosome inhibitors, including Madrasin and Isoginkgetin, also produced robust Z-NA signals similar to those of PladB (Figure S2A, B). Collectively, these results indicate that spliceosome inhibition caused the accumulation of endogenous Z-form RNA in cells.

**Figure 3.**
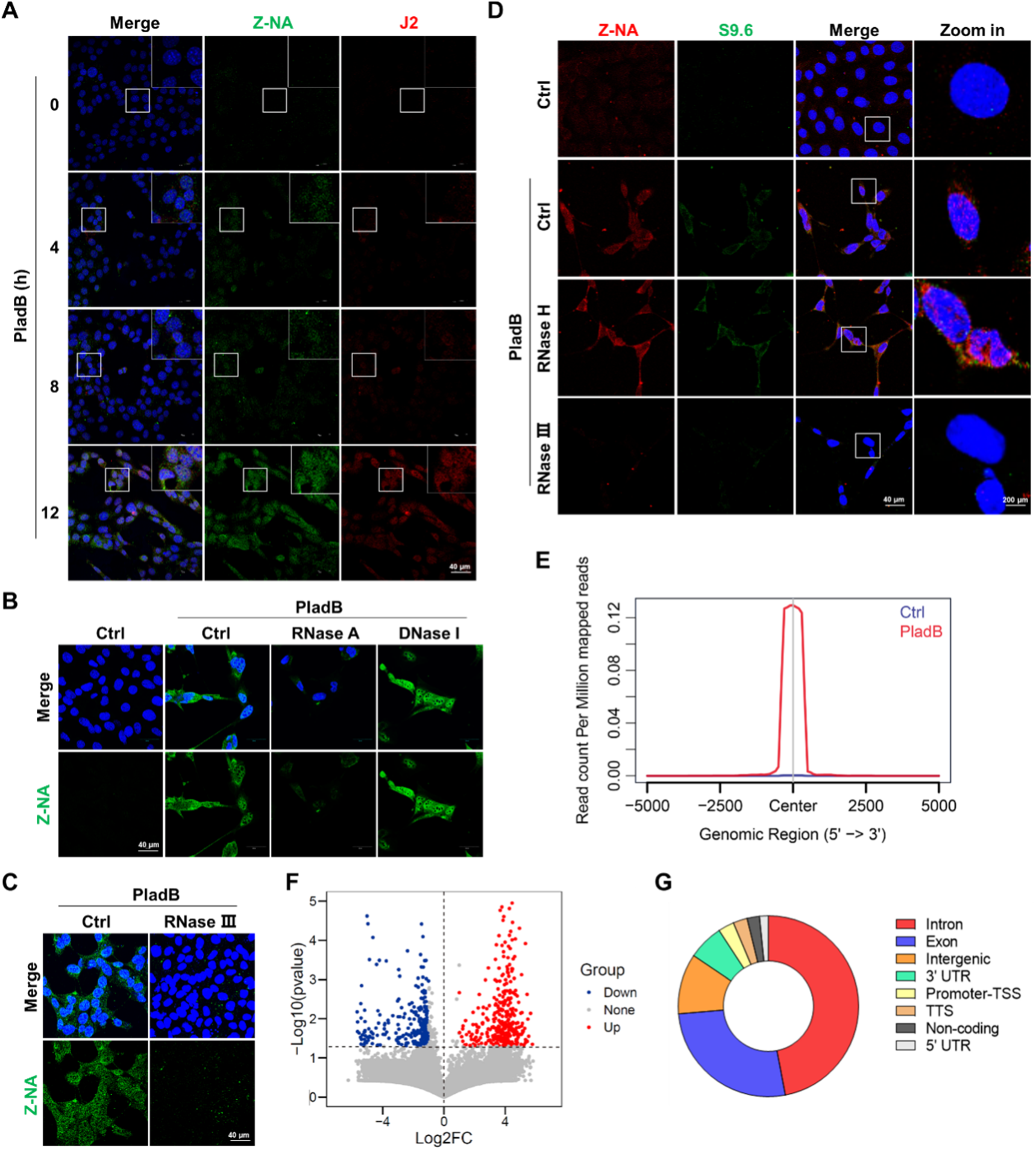
Inhibition of spliceosome caused accumulation of Z-NA. **(A)** Immunofluorescence staining of Z-NA (green) and dsRNA (J2, red) in MEFs treated with PladB (100 nM) as indicated. **(B)** MEFs treated with PladB for 24 h were exposed to the indicated nucleases for 60 min and stained for Z-NA. **(C)** MEFs treated with PladB for 24 h were exposed to RNase III for 60 min and stained for Z-NA. **(D)** MEFs treated with PladB for 24 h were exposed to the indicated nucleases for 60 min and stained for Z-NA and S9.6. **(E)** Metaplot of RIP-RNA signals in the significantly up- and down-regulated peak center (n = 584). **(F)** Volcano plot showing PladB-induced peaks in MEFs. Red and blue dots denote PladB-upregulated (n = 320) and downregulated peaks (n = 264), respectively. **(G)** Pie charts showing the distributions of upregulated peaks identified by RIP-seq of PladB treated MEFs (n = 320).

We then sought to investigate the source of the Z-RNA in response to spliceosome inhibition. Spliceosome perturbations induce transcriptome-wide defects in RNA splicing, including intron retention, but the characteristics of these intron-retained transcripts is unclear. To directly assess the composition of Z-RNA that recognized by ZBP1 after spliceosome inhibition, we utilized the ZBP1 antibody to immunoprecipitate Z-RNAs from these cells following PladB treatment. Sequencing ZBP1-enriched RNAs revealed that most (> 70%) of the RNAs were intronic RNAs and exonic RNAs from protein-coding mRNAs; intergenic, UTR, noncoding RNA and other non-mRNA species constituted the remainder of the reads (Figure 3E-G). Although these Z-RNA sequences were mapped to various genes involving diverse biological processes and pathways (Figure S3D and E), most of them harbored a highly consensus motif (Figure S3F).

### Spliceosome inhibition induced the egress of Z-RNA from nucleus to cytoplasm for ZBP1 sensing

Although ZBP1 is primarily located in the cytoplasm, it can also shuttle between the nucleus and cytoplasm. To test whether ZBP1 sensed Z-RNAs in the nucleus or in the cytoplasm, we first examined the cellular location of ZBP1 following PladB treatment by stably reconstituting *Zbp1* KO MEFs with YFP-tagged ZBP1. As expected, we observed that ZBP1 primarily located in the cytoplasm, with weak signals in the nucleus in untreated MEFs (Figure 4A). PladB-treatment did not significantly alter the pattern of ZBP1 in the cell until the late stage, when obvious cell death occurred, suggesting that ZBP1 would not initiates necroptosis from the nucleus (Figure 4A). Immunoblot analysis of the cytoplasmic and nuclear extracts from cells that were stimulated with PladB for a series of time revealed that the ZBP1 proteins were only found in the cytosolic fraction (Figure 4B). Notably, the necroptosis signal, pMLKL started at approximately 4 h post-treatment; these signals were observed only in the cytoplasm but not the nucleus and becoming strong in the cytoplasm around 12 h post-treatment (Figure 4B). These results indicate that ZBP1 sensed Z-RNAs and activates RIPK3-MLKL necroptosis signaling in the cytoplasm.

**Figure 4.**
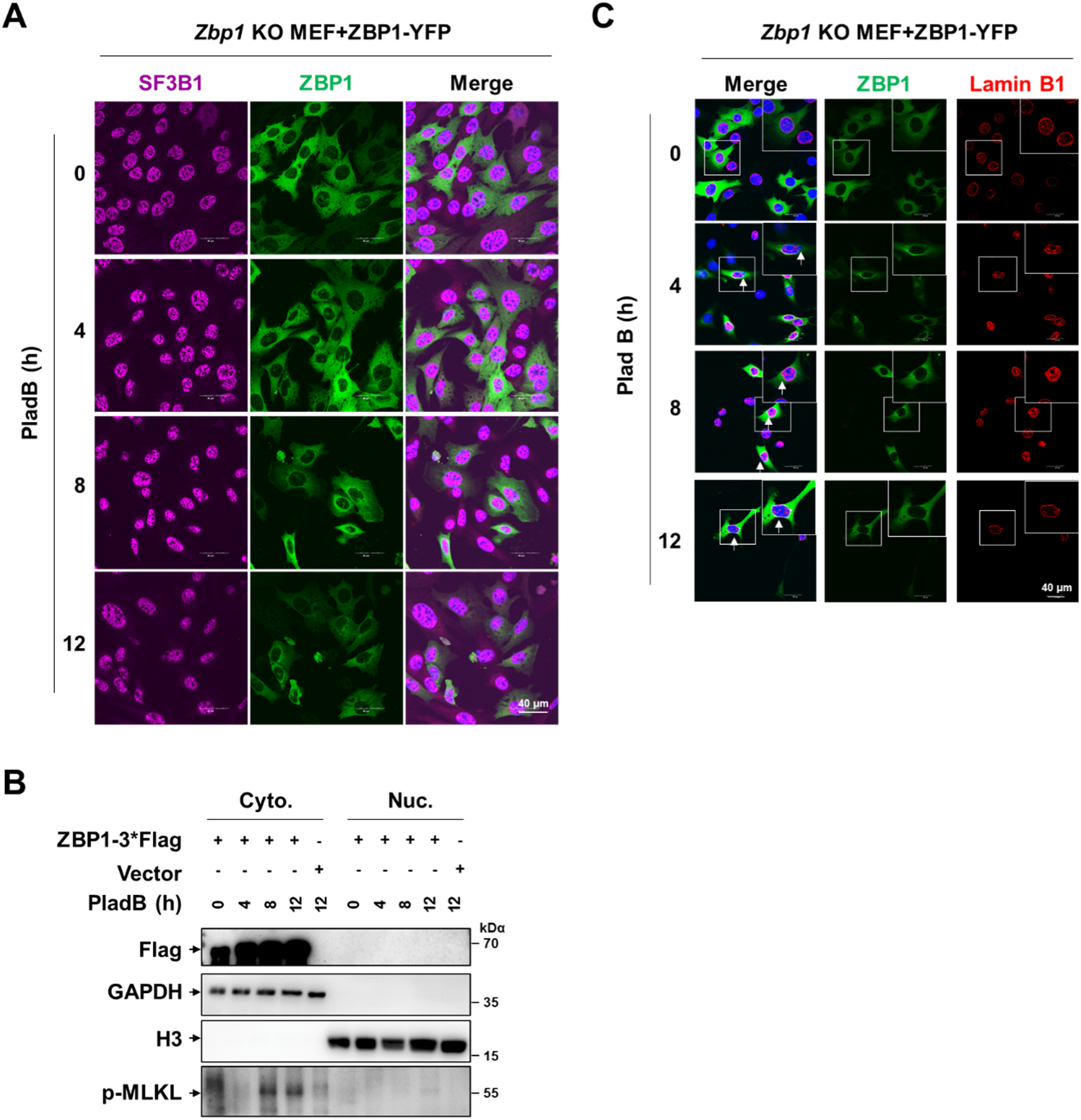
Spliceosome inhibition induced the egress of Z-RNA from nucleus to cytoplasm for ZBP1 sensing. **(A)** Immunofluorescence staining for SF3B1 (magenta) in YFP–ZBP1 reconstituted MEFs treated with PladB as indicated. **(B)** Immunoblot analysis of p-MLKL in nuclear (Nuc.) and cytoplasmic (Cyto.) fractions from ZBP1-3*Flag reconstituted MEF cells treated with PladB as indicated. **(C)** Immunofluorescence staining for lamin B1 in YFP–ZBP1 reconstituted MEFs treated with PladB as indicated.

RNA splicing occurs in the nucleus. As the Z-RNA signals were first detected mainly in the nucleus and late throughout the whole cells along with the sensing by ZBP1 initiated in the cytoplasm, we reasoned that the Z-RNA produced by spliceosome inhibition may egress from the nucleus to the cytoplasm. By staining the nuclear lamina protein lamin B1, we observed abnormalities of the nuclear envelope at the early stage of PladB-treatment (Figure 4C). The nuclear envelope, while distorted, invaginated, is rarely breached, and DNA leakage into the cytosol is not typically observed (Figure 4C), suggesting that the Z-RNA may not simply released into the cytoplasm through breakdown of the nuclear envelope. It has been reported that the phospholipids within both the inner and outer nuclear membranes might draw activated MLKL to nuclear membranes and lead to their rupture during IAV-activated necroptosis and defective prelaminin A-promoted necroptosis^35, 36^. Of note, the distortion of the nuclear envelope as well as the dispersal of Z-RNA upon PladB treatment were independent of the activated MLKL, as deletion of *Mlkl* has no impact on this change (Figure S3A), which is in consistent with the observation that no pMLKL could be observed in the nuclear fraction. In addition, the abnormality of the nuclear envelops induced by PladB treatment still occurred even the cell death were totally blocked by deleting both *Ripk3* and *Casp8* (Figure S3B). These results suggest that the egress of Z-RNA from the nucleus to the cytoplasm is an active process involving nuclear export, independent of cell death process. The detail mechanism by which Z-RNA exits the nucleus required further investigation.

## Discussion

Our findings suggest that the activation of necroptosis through Z-NA sensing by ZBP1 may represent a common mechanism that responds to widespread RNA splicing aberrations across cancer and other disease states (Figure S4). While well-established pathways like the DNA damage response (DDR) and the unfolded protein response (UPR) handle macromolecular issues such as DNA and protein anomalies, coordinating responses to restore cellular balance or induce cell death, there has been less clarity about how cells respond to widespread RNA splicing errors^33, 37, 38^. Although quality-control mechanisms like nonsense-mediated decay (NMD) exist for RNA, it has been unclear whether there are signaling pathways that sense widespread mis-splicing of RNA to dictate cell fate.

Our evidence indicates that Z-RNA sensing and the resulting ZBP1-dependent cell death might be part of a coordinated response to widespread RNA splicing aberrations. The mechanism of how Z-RNAs are exported from the nucleus and accumulate in the cytoplasm remains unclear. The different RNA species that are produced in the nucleus are exported through the nuclear pore complexes via mobile export receptors. Among these, the nuclear transport receptor exportin-1, which is involved in the export of nucleic acids, had no role in regulating the egress of the Z-NA from the nucleus to the cytoplasm induced by PladB treatment (Figure S3C). Notably, SF3B1 mutants have been linked to altered RNA export, suggesting that spliceosome inhibition might disrupt RNA export processes. Overall, our results highlight Z-RNA-mediated necroptosis as a potential mechanism for sensing and responding to broad cellular splicing dysfunction (Figure S4).

Spliceosome inhibitors have been shown to be well-tolerated in human Phase 1 clinical trials, but the clinical outcomes are not favorable^39^. This may be due to low-to-absent basal RIPK3 levels in many cancer cell types, making them resistant to ZBP1-mediated necroptosis. ZBP1 expression is, however, strongly induced by IFN. The chronic IFN signature in many tumors may upregulate ZBP1 expression selectively in cells of the tumor microenvironment (TME), creating a potential therapeutic opportunity for systemic administration of spliceosome inhibitors. It’s also important to note that while RIPK3 expression is typically silenced in tumor cells, immune cells such as macrophages, T cells, B cells, and neutrophils have relatively high levels of RIP3. These immune cells might be particularly sensitive to spliceosome inhibitors, and their death could impact anti-tumor immune responses. Investigating these aspects will be crucial in determining whether spliceosome-targeted therapies can effectively kill tumor cells or enhance immune responses against aggressive, immune-cold tumors.

## Acknowledgements

The study was supported by the National Natural Science Foundation of China (32225016 to W.M., 32170751 to Z.-H.Y.), the National Key R&D Program of China (2021YFA1101401 to W.M.), the Leading Innovation and Entrepreneurship Team of Zhejiang Province (2023R01005 to W.M. and Z.-H.Y.).

## Author contributions

Conceptualization and Supervision: Z.-H.Y., Z.-Y.C. and W.M.; Data curation: Z.-H.Y., P.W. and Z.-Y.C. with the help from C.-R.Y., J.H., L.W. and S.-H.G.; RIP-seq analysis: Y.Z., Y.M. and Y.Y.; Formal analysis: Z.-H.Y., P.W., R.Z., H.S., Z.-Y.C. and W.M.; Writing-original draft: Z.-H.Y., Z.-Y.C. and W.M.; Writing-review & editing: Z.-H.Y. and W.M.

## STAR★METHODS

Detailed methods are provided in the online version of this paper and include the following:

- KEY RESOURCES TABLE
- RESOURCE AVAILABILITY

○ Lead contact
○ Materials availability
○ Data and code availability
- EXPERIMENTAL MODEL AND SUBJECT DETAILS

○ Cell lines
- METHOD DETAILS

○ Generation of knockout cell lines
○ Cell death assay
○ Plasmid construction
○ Lentivirus preparation and infection
○ Subcellular fractionation
○ Western blot
○ Quantitative PCR
○ Immunofluorescence staining and imaging
○ Cross-link immunoprecipitation
○ RIP-seq analysis
- QUANTIFICATION AND STATISTICAL ANALYSIS

### KEY RESOURCES TABLE

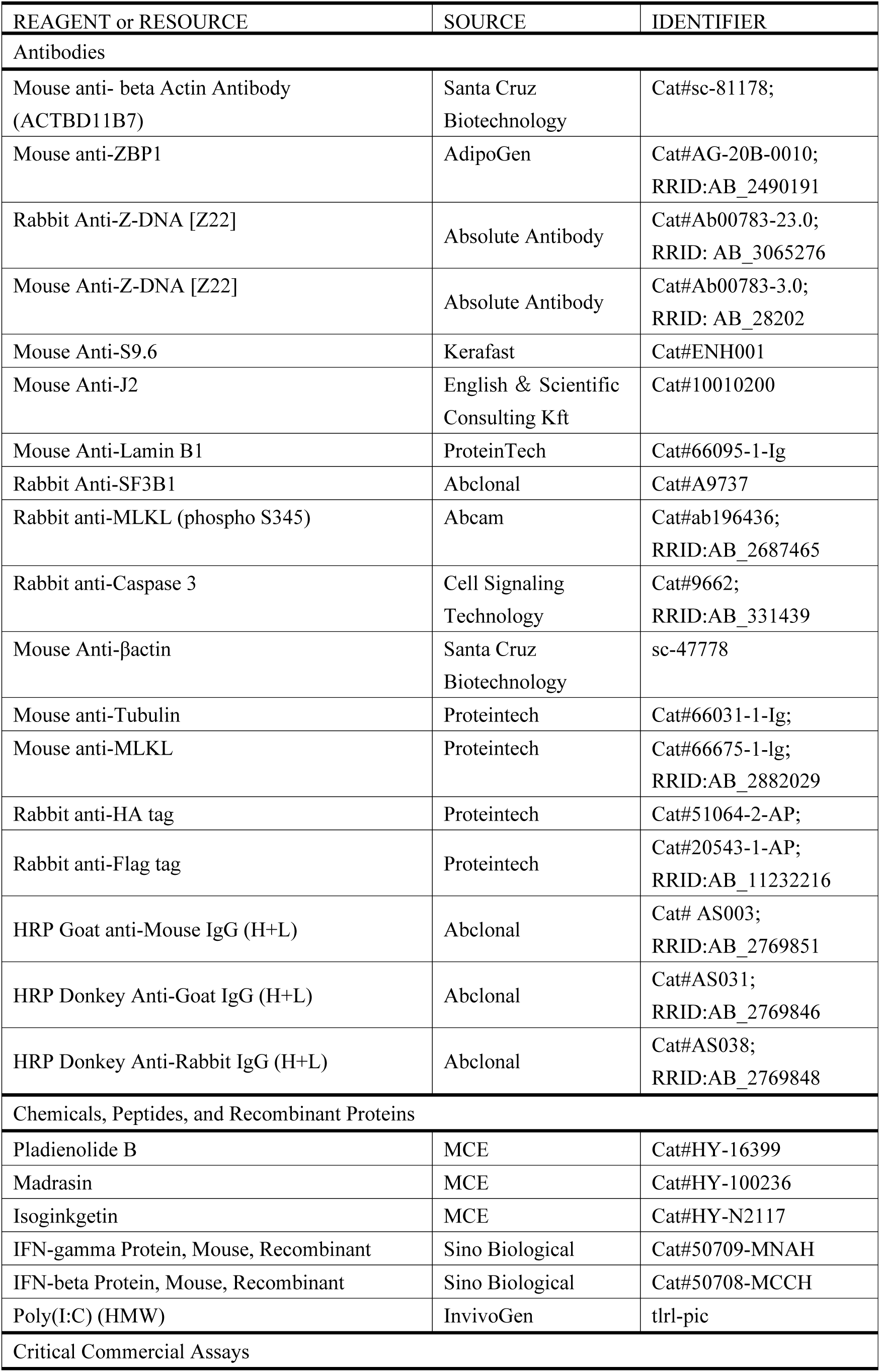
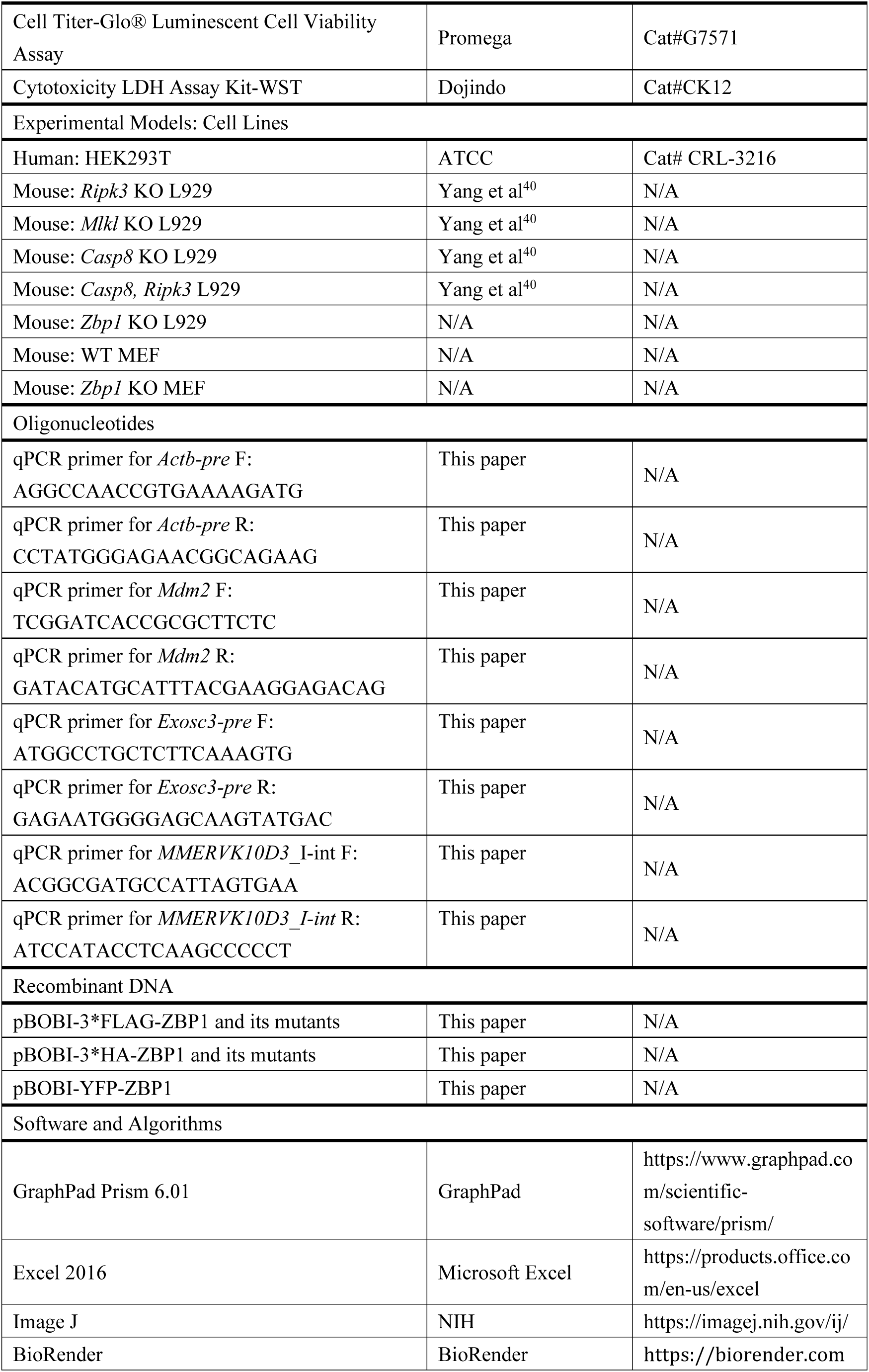
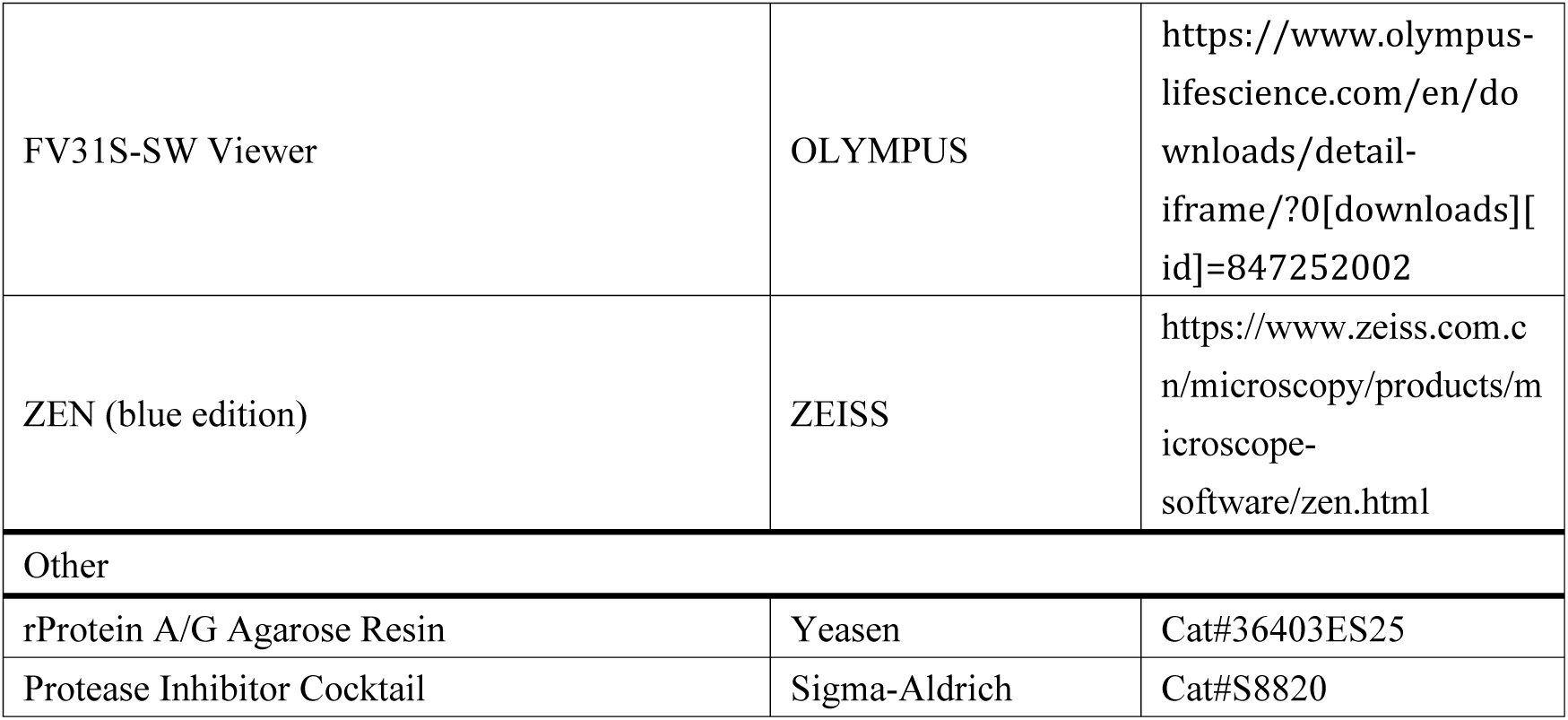

### RESOURCE AVAILABILTY

#### Lead contact

Further information and requests for resources and reagents should be directed to and will be fulfilled by the Lead contact, Wei Mo.

#### Materials availability

All plasmids, reagents and cell lines generated in this study are available from the Lead contact.

#### Data and code availability

- Immunoblot images data and microscopy images reported in this paper will be shared by the Lead contact upon request.
- This paper does not report original code. All source data for RIP-seq have been deposited at Gene Expression Omnibus under accession code GSE276187 and are publicly available as of the date of publication.
- Any additional information required to reanalyze the data reported in this paper is available from the Lead contact upon request.

### EXPERIMENTAL MODEL AND SUBJECT DETAILS

#### Cell lines

HEK293T cells, Mouse fibrosarcoma WT L929, *Tnfr1* KO, *Casp8* KO, *Mlkl* KO, *Ripk3* KO, *Casp8, Ripk3* DKO L929 cells were kindly provided by Prof. Jiahuai Han. MEF was generated and infected as described before^40^. Briefly, All MEFs were from E13.5 embryos and infected with the retroviral Large T. Cells were cultured in Dulbecco′s Modified Eagle′s Medium (Gibco, USA) supplemented with 10% FBS (vol/vol) (HyClone, USA), 100 IU penicillin (Sangon, China) and 100 mg/ml streptomycin (Sangon, China) at 37 °C in a humidified incubator containing 5% CO_2_. All cell lines were well established and frequently checked by monitoring morphology and functionalities. All the cell lines were authenticated by STR analysis and were routinely tested to be mycoplasma-free.

### METHOD DETAILS

#### Generation of knockout cell lines

*Zbp1* knockout (KO) L929 and MEF cell lines were generated using CRISPR/Cas9 methods. The target sequence for mouse *Zbp1* were: 5’-caggtgttgagcgatgacgg-3’. gRNA was transduced into indicated cell line by lentiviral delivery. Cells were then subjected to blasticidin selection. Single-cell clones were isolated from the selected pool by limiting dilution cloning in 96-well plates and then screened for indicated gene expression by western blot. Selected knockout clones were further verified by DNA sequencing.

#### Cell death assay

Cell death was analyzed by using the Cytotoxicity LDH Assay kit-WST (Dojindo Molecular Technologies) according to the manufacturer’s instructions. Briefly, 2 × 10^4 cells were seeded in 96-well plates with white walls (Nunc). After 12 h, the cells were treated with reagents for the indicated durations. After treatment, an equal volume of medium was used for measurement of LDH release.

#### Plasmid construction

The full-length sequences of ZBP1 were amplified from cDNA. ZBP1 mutations were introduced by two-round PCR. All these DNA fragments were cloned into pBOBI lentiviral vectors with Flag/HA/YFP tags. The Exo III-assisted ligation-independent cloning method was used for subcloning. All plasmids were verified by DNA sequencing.

#### Lentivirus preparation and infection

HEK293T cells were transfected with lentiviral vectors carrying cDNAs or gRNA of interest and lentivirus-packing plasmids (PMD2/PSPAX) by the calcium phosphate precipitation method. 12 hours later, cell culture medium was changed and the virus-containing medium was collected 36 hours later and used for infection. For infection, target cells were infected with lentivirus containing medium in the presence of 10 µg/ml Polybrene and then centrifuged at 2,500 rpm for 30 min. The medium was changed 12 h later.

#### Subcellular fractionation

The cells were lysed in hypotonic buffer (20 mM Tris-HCl, pH 7.4, 10 mM NaCl, 3 mM MgCl_2_) containing 0.5% NP-40 at 4°C for 10 minutes. The lysate was centrifuged at 4°C at 3000 rpm for 10 minutes to separate the cytosolic and nuclear fractions. The nuclear fraction was lysed using RIPA buffer (50 mM Tris-HCl, pH 7.4, 1% NP-40, 12 mM Sodium deoxycholate, 3.5 mM Sodium dodecyl sulfate, 150 mM NaCl and 2 mM EDTA), followed by moderate sonication. Both the cytosolic fraction and the RIPA-lysed nuclear component were centrifuged again at 4°C at 15000 rpm for 15 minutes.

#### Western blot

Cell lysates were lysed in RIPA buffer (50 mM Tris-HCl, pH7.4, 1% NP-40, 12 mM Sodium deoxycholate, 3.5 mM Sodium dodecyl sulfate, 150 mM Sodium chloride and 2 mM EDTA) supplemented with protease inhibitor cocktail (MCE). Cell lysates were incubated on ice for 10 min, and briefly sonicated to shear chromatin, then cleared by high speed centrifugation (12,000 g, 30 min) at 4℃. The supernatants were used for analysis. For analysis of phosphorylated MLKL, the insoluble fractions were dissolved in 1× SDS loading buffer for western blot. The protein samples were resolved by SDS-PAGE, transferred onto a polyvinylidene difluoride (PVDF) membrane, and probed with corresponding antibodies. The antibodies used were listed in the Key Resource Table.

#### Quantitative PCR

Total RNA was extracted from cells using Trizol reagent according to the manufacturer’s instructions, and reverse transcribed into cDNA with the GoScript Reverse Transcription System (Promega). qPCR (ChamQ Universal SYBR qPCR Master Mix, Vazyme) was performed using the CFX Connect Real-Time PCR Detection System (Bio-Rad). Detailed information about the primers sequences is listed in Key Resource Table.

#### Immunofluorescence staining and imaging

After stimulation, cells were washed twice with PBS followed by fixation for 15 min at room temperature in freshly prepared 4% paraformaldehyde. The fixed cells were then permeabilized in 0.5% Triton X-100/PBS and non-specific binding was blocked with 3% BSA in PBS. Cells were incubated with the following antibodies for X hours at room temperature: anti-pMLKL (1:1000, Abcam, ab196436), anti-Lamin B1 (1:1000, ProteinTech, 12987-1-AP), anti-ZNA (1:1000, Absolute Antibody, Ab00783-23.0), anti-J2 (1:1000, SCICONS, 10010200). The secondary antibodies used were Alexa Fluor 568-conjugated anti-rabbit IgG (Life Technologies, A11036; 1:500), Alexa Fluor 488-conjugated anti-rabbit IgG (Life Technologies, A11034; 1:500), Alexa Fluor 488-conjugated anti-mouse IgG (Life Technologies, A11029; 1:500), Alexa Fluor 647-conjugated anti-mouse IgG (Life Technologies, A21236; 1:500). Cells were counterstained with Hoechst to visualize nuclei. For enzymatic digestion, RNase A (1 mg/ml) or DNase I (25 U/ ml) was added for 1h at 37°C before primary antibody incubation. All images were acquired on a Olympus 20040551 laser scanning confocal microscope using a 40×objective or 60×objective. Unprocessed images were analyzed by ImageJ software.

#### Cross-link immunoprecipitation

Cross-link immunoprecipitation (CLIP) of MEF cells were referred to Wang et al^22^. Briefly, cells were overlaid with 1.5 mL PBS and crosslinked with 150 mJ/cm^2^ (254 nm) (CX-2000, analytic Jena). Cells were lysed in 1 ml RIPA buffer (50 mM Tris-HCl pH 8.0, 150 mM NaCl, 1% NP-40, 0.1% SDS, 0.5% deoxycholate, 2 mM EDTA) for 30 min, and then 12.5 μL RNasin Plus (Promega) was added, followed by centrifuging at 12,000 rpm for 15 min. Samples were incubated with 2 μg of anti-ZBP1 antibodies overnight at 4℃. After immunoprecipitation with Protein A/G agarose beads (Millipore), the beads were beads were washed twice with high-salt wash buffer (50 mM Tris-HCl pH 8.0, 500 mM NaCl, 1% NP-40, 0.1% SDS, 0.5% deoxycholate) and once with RIPA buffer. After wash, beads were resuspended in 100 μl of reverse buffer (100 mM Tris-HCl pH8.0, 10 mM EDTA, 1% SDS, 100 mM DTT) with 10 μl 5M NaCl and 50 μg proteinase K. Beads were then incubated in a heating block at 42 °C for 1 h to digest peptide, followed by 65 °C for 1.5 h to reverse the crosslink. The immunoprecipitated RNA was extracted with the TRIzol method, and purified RNA was used for RNA sequencing. Libraries were quantified using a Quant-iT PicoGreen ds DNA assay (Thermo Fisher Scientific) or by low-pass sequencing with a MiSeq nano kit (Illumina). Paired-end 150 cycle sequencing was performed on a NovaSeq 6000 (Illumina).

#### RIP-seq analysis

For RIP-seq analysis, the adapter sequences of raw reads were initially removed using cutadapt v4.0. Subsequently, the processed reads were aligned to the mouse genome mm10 using Hisat2 v2.2.1 with default parameters. Peak calling was done using macs2 v2.2.7.1 with these key parameters: -f BED –nolambda --nomodel --extsize 50. Peak distribution across genomic regions was calculated using HOMER v4.11 annotatePeaks.pl. Differential peaks were defined as peaks with *p* value less than 0.05 and fold change larger than 2. The following Pathways and GO analysis were annotated in the KEGG (Kyoto Encyclopedia of Genes and Genomes) database phyper and shown by ggplot2 (https://ggplot2.tidyverse.org/). Motif analysis was performed using MEME (https://meme-suite.org/meme/tools/meme).

### QUANTIFICATION AND STATISTICAL ANALYSIS

No statistical methods were used to predetermine sample size. Statistical analysis was performed with Prism software (GraphPad Software). Data are presented as the means ± SD. A two-tailed Student’s *t*-test was used to compare differences between treated groups and their paired controls. Differences in compared groups were considered statistically significantly different with P values: ns: P≥0.0.5; *: P <0.05; ** : P <0.01; ***: P <0.001; ****: P< 0.0001.

---

**Figure S1.**
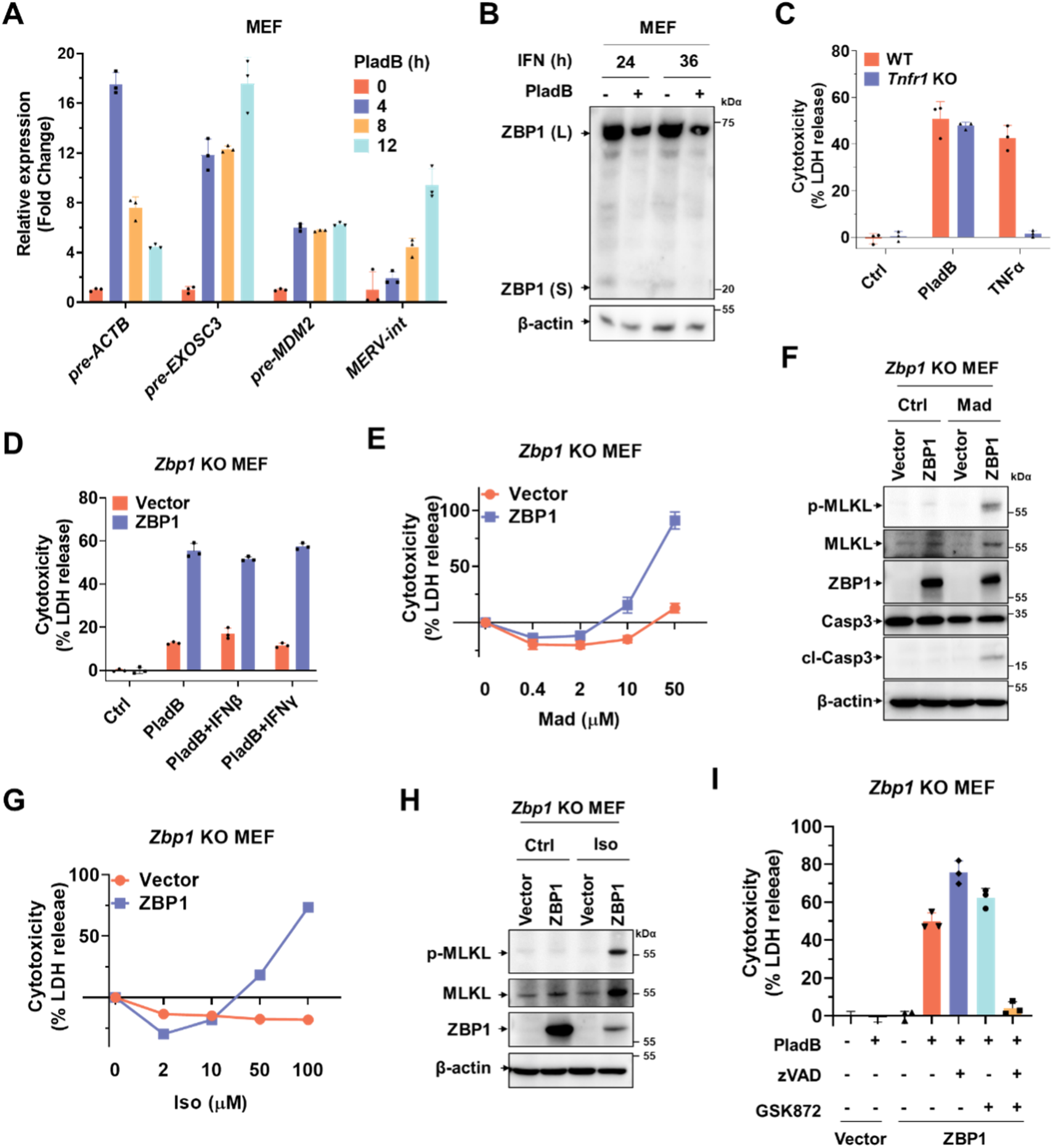
Inhibition of spliceosome initiates ZBP1-dependent cell death. **(A)** qPCR analysis of pre-mRNA level in MEFs treated with PladB as indicated. **(B)** Immunoblot analysis of WT MEFs were treated with PladB plus IFNγ (prime for 24 h or 36 h) after 12 h. **(C)** IFN-priming WT, *Tnfr1* KO L929 cells were treated with PladB or TNFα (10 ng/ml) for 24 h and the cell death was measured by LDH release. **(D)** *Zbp1* KO MEF reconstituted with empty vector or full-length murine ZBP1 were treated as indicated for 24 h and the cell death was measured by LDH release. **(E)** *Zbp1* KO MEFs reconstituted with empty vector or full-length murine ZBP1 treated with Madarsin (Mad) as indicated for 12 h and the cell death was measured by LDH release. **(F)** The same cells as in (**E**) were treated with Mad (50 μM) as indicated for 12 h and the cell lysates were analyzed by immunoblot. **(G)** *Zbp1* KO MEFs reconstituted with empty vector or full-length murine ZBP1 were treated with Iso as indicated for 12 h and the cell death was measured by LDH release. **(H)** The same cells as in (**G**) were treated with Iso (50 μM) as indicated for 12 h and the cell lysates were analyzed by immunoblot. **(I)** *Zbp1* KO MEFs reconstituted with empty vector or full-length murine ZBP1 were treated with PladB, in the presence of GSK’872 (10 μM), zVAD (50 μM), or both inhibitors together as indicated for 12 h and the cell death was measured by LDH release. Data are representative of three independent experiments. Error bars represent mean ± SD. *p<0.05, **p<0.01, ***p<0.001, ****p<0.0001.

**Figure S2.**
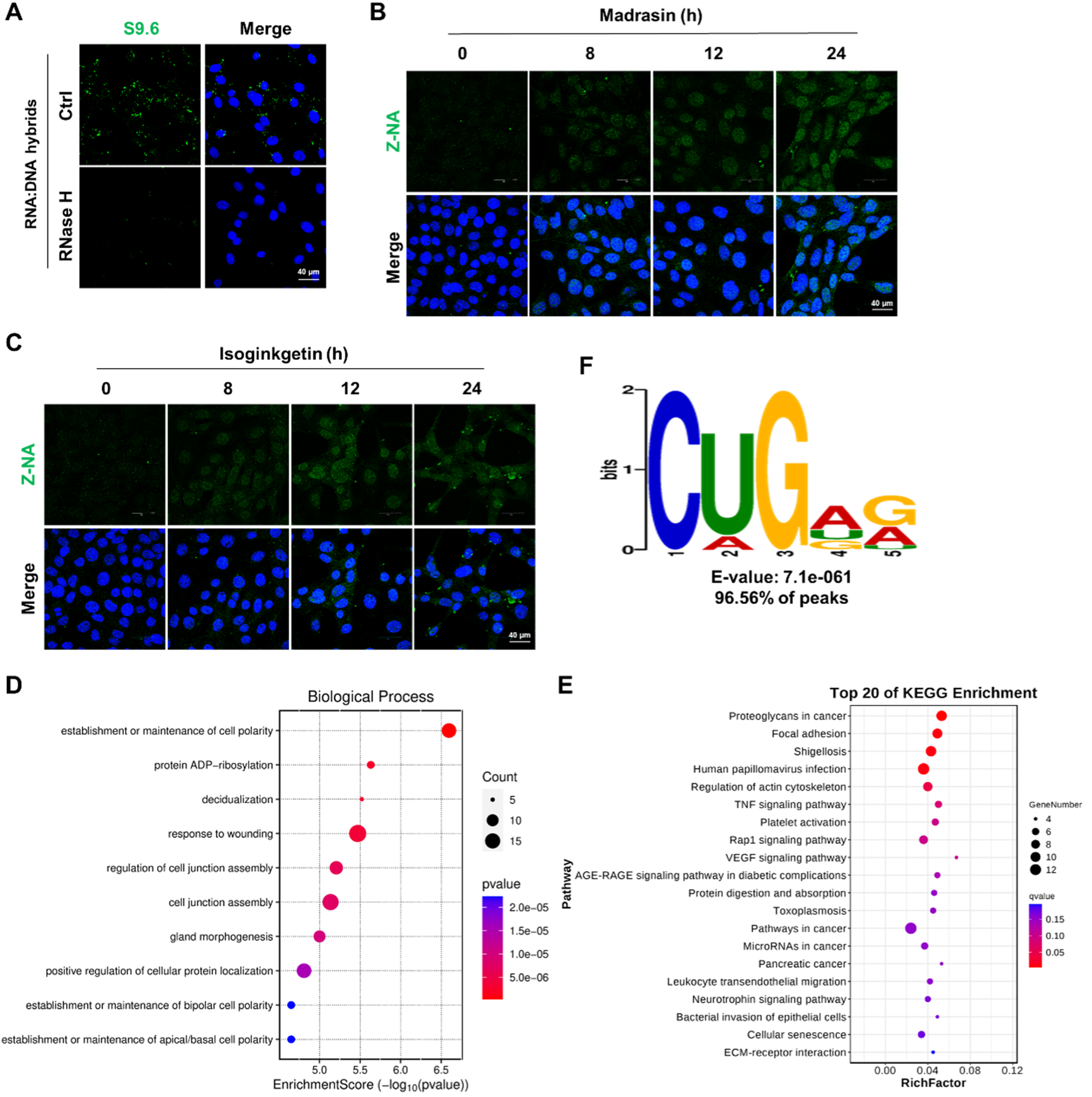
Inhibition of spliceosome caused accumulation of Z-NA. **(A)** MEFs transfected with RNA:DNA hybrids were exposed to the indicated nucleases for 60 min and stained with S9.6 antibody. **(B)** Immunofluorescence staining of Z-NA (green) in MEFs treated with Madrasin as indicated. **(C)** Immunofluorescence staining of Z-NA (green) in MEFs treated with Isoginkgetin as indicated. **(D)** GO analysis of PladB-upregulated genes in ZBP1 RIP-seq. **(E)** Bar plots depicting the *p* values of relevant pathways related to PladB-upregulated genes in ZBP1 RIP-seq. **(F)** The motif with the most significant *p* value according to the HOMER analysis is displayed.

**Figure S3.**
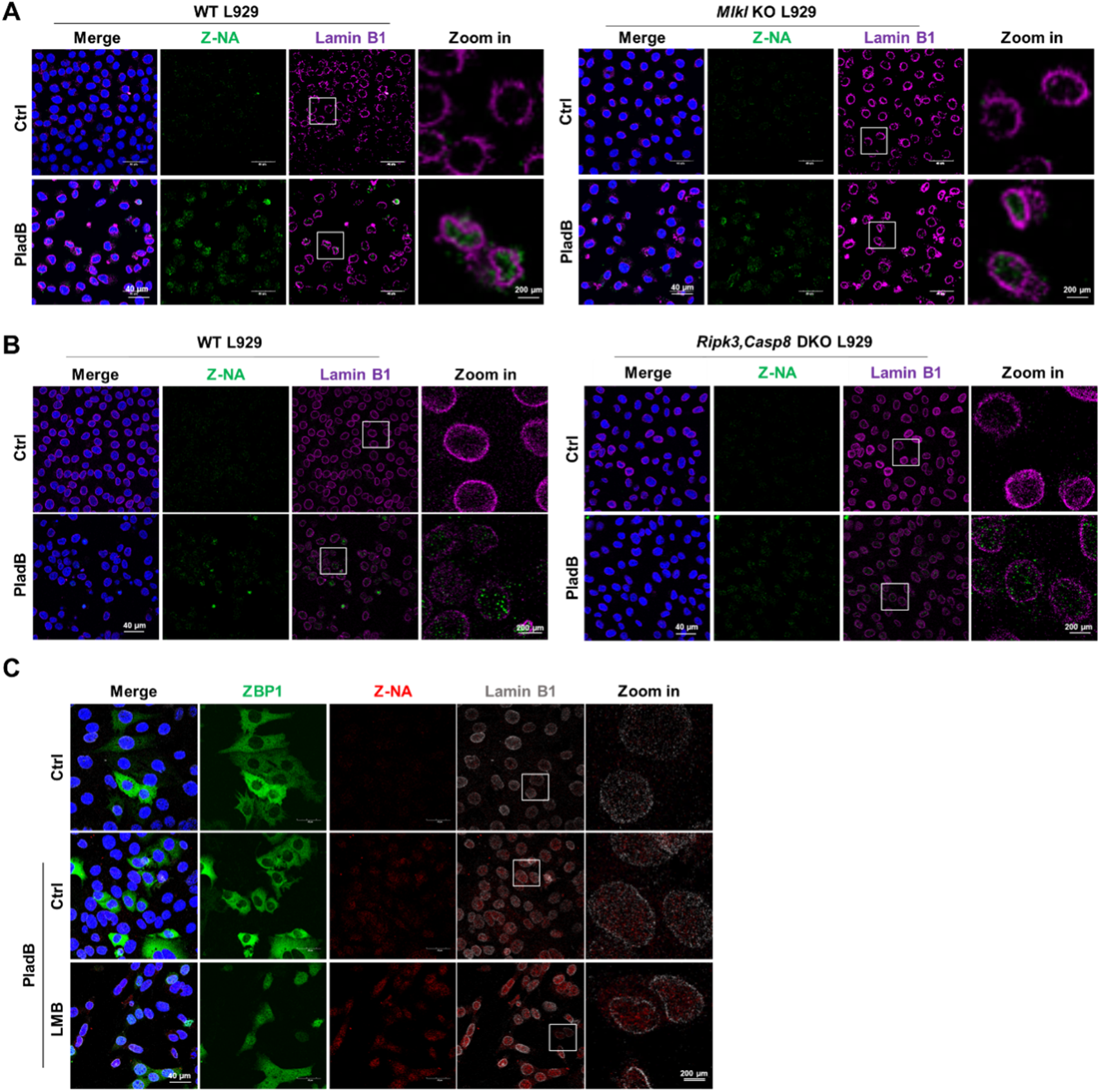
Spliceosome inhibition induced the egress of Z-RNA from nucleus to cytoplasm independent of cell death and Exportin-1. **(A)** Immunofluorescence staining of Z-NA (green) and Lamin B1 (magenta) in IFN-primed WT or *Mlkl* KO L929 cells treated with PladB for 24 h. **(B)** Immunofluorescence staining of Z-NA (green) and Lamin B1 (magenta) in IFN-primed WT or *Ripk3&Casp8* DKO L929 cells treated with PladB for 24 h. **(C)** Immunofluorescence staining for Z-NA (red) and Lamin B1 (gray) in YFP–ZBP1 reconstituted MEFs treated with PladB or PladB plus LMB (5 ng/ml) for 12 h as indicated.

**Figure S4.**
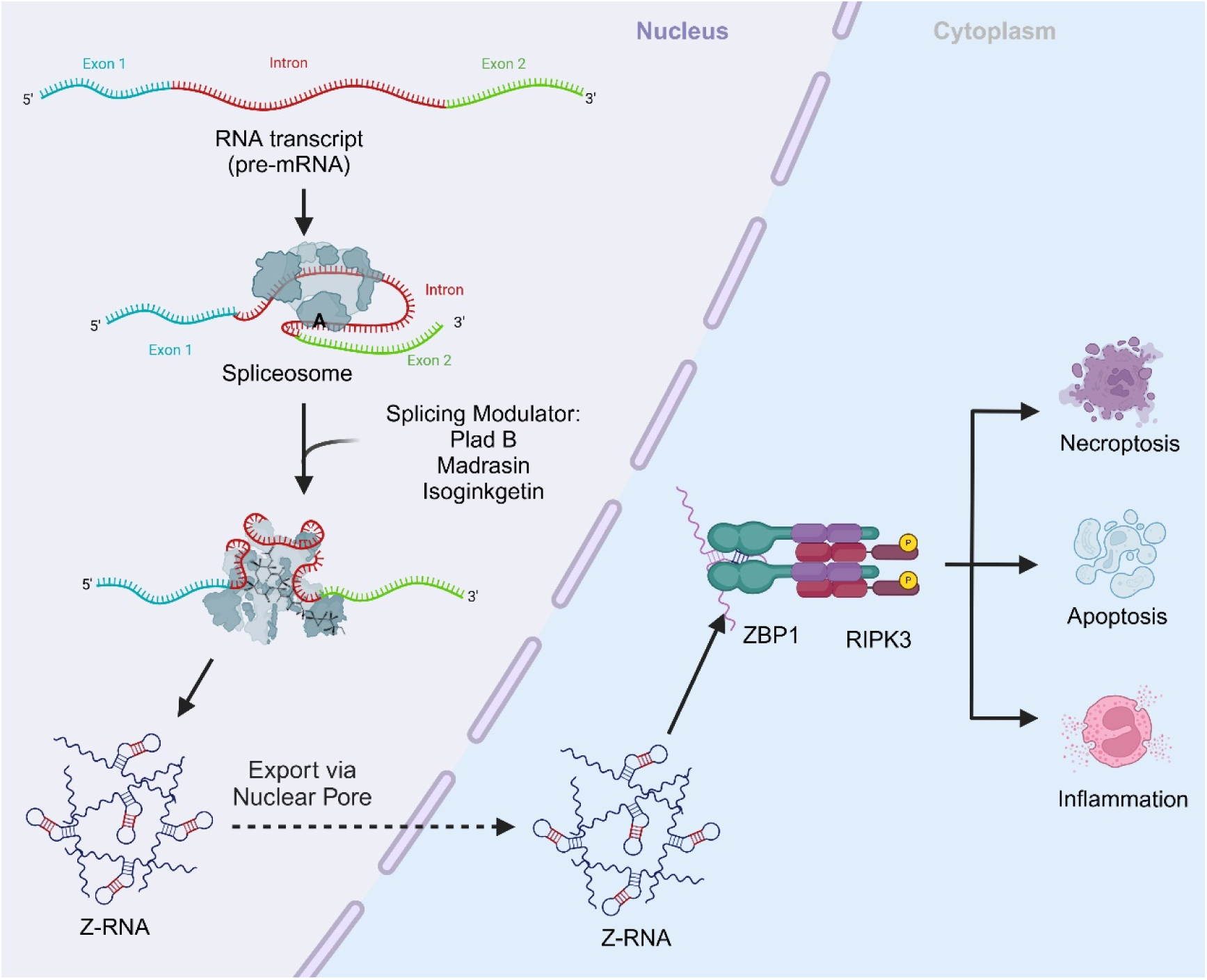
ZBP1 Senses Splicing Aberration through Z-RNA to promote Cell Death. Inhibition of spliceosome caused he accumulation of Z-form dsRNA (Z-RNA) in the nucleus. These Z-RNAs exit from the nucleus to the cytoplasm by active nuclear export machinery. Cytosolic ZBP1 sensed these danger-associated molecular pattern (DAMP) and induced cell death, including necroptosis and apoptosis. Our findings suggest that the activation of cell death through Z-RNA sensing by ZBP1 may represent a common mechanism that responds to widespread RNA splicing aberrations.

